# Adaptive Estimation for Epidemic Renewal and Phylogenetic Skyline Models

**DOI:** 10.1101/703751

**Authors:** Kris V Parag, Christl A Donnelly

## Abstract

Estimating temporal changes in a target population from phylogenetic or count data is an important problem in ecology and epidemiology. Reliable estimates can provide key insights into the climatic and biological drivers influencing the diversity or structure of that population and evidence hypotheses concerning its future growth or decline. In infectious disease applications, the individuals infected across an epidemic form the target population. The renewal model estimates the effective reproduction number, *R*, of the epidemic from counts of its observed cases. The skyline model infers the effective population size, *N*, underlying a phylogeny of sequences sampled from that epidemic. Practically, *R* measures ongoing epidemic growth while *N* informs on historical caseload. While both models solve distinct problems, the reliability of their estimates depends on *p*-dimensional piecewise-constant functions. If *p* is misspecified, the model might underfit significant changes or overfit noise and promote a spurious understanding of the epidemic, which might misguide intervention policies or misinform forecasts. Surprisingly, no transparent yet principled approach for optimising *p* exists. Usually, *p* is heuristically set, or obscurely controlled via complex algorithms. We present a computable and interpretable *p*-selection method based on the minimum description length (MDL) formalism of information theory. Unlike many standard model selection techniques, MDL accounts for the additional statistical complexity induced by how parameters interact. As a result, our method optimises *p* so that *R* and *N* estimates properly adapt to the available data. It also outperforms comparable Akaike and Bayesian information criteria on several classification problems. Our approach requires some knowledge of the parameter space and exposes the similarities between renewal and skyline models.

## I. Introduction

Inferring the temporal trends or dynamics of a target population is an important problem in ecology, evolution and systematics. Reliable estimates of the demographic changes underlying empirical data sampled from an animal or human population, for example, can corroborate or refute hypotheses about the historical and ongoing influence of environmental or anthropogenic factors, or inform on the major forces shaping the diversity and structure of that population [40] [13]. In infectious disease epidemiology, where the target population is often the number of infected individuals (infecteds), demographic fluctuations can provide insight into key shifts in the fitness and transmissibility of a pathogen and motivate or validate public health intervention policy [34] [3].

Sampled phylogenies (or genealogies) and incidence curves (or epi-curves) are two related but distinct types of empirical data that inform about the population dynamics and ecology of infectious epidemics. Phylogenies map the tree of ancestral relationships among genetic sequences that were sampled from the infected population [6]. They facilitate a retrospective view of epidemic dynamics by allowing estimation of the historical effective size or diversity of that population. Incidence curves chart the number of infecteds observed longitudinally across the epidemic [43]. They provide insight into the ongoing rate of spread of that epidemic, by enabling the inference of its effective reproduction number. Minimal examples of each empirical data type are given in (a)(i) and (b)(ii) of Fig. 1.

**Fig. 1:**
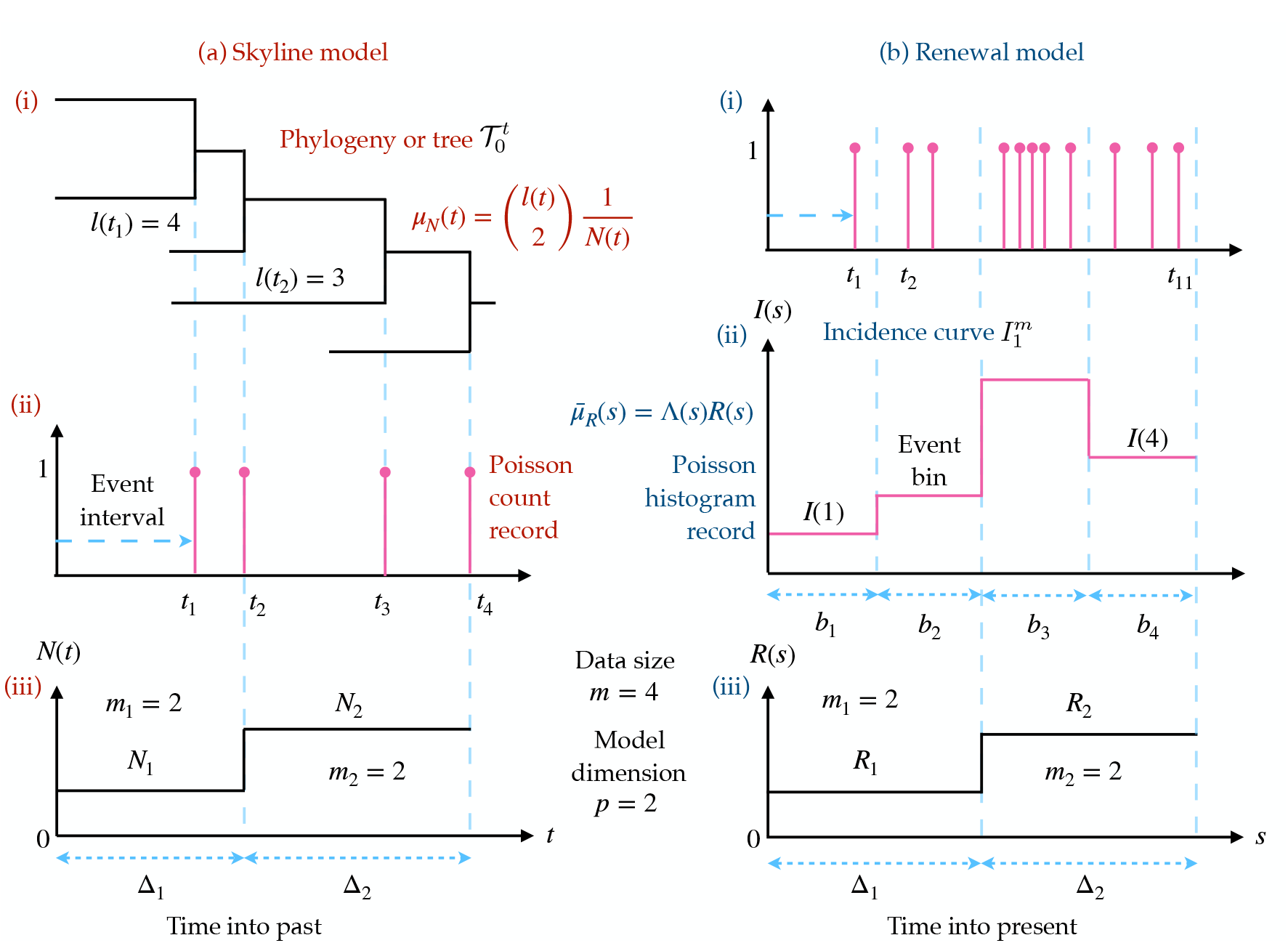
Skyline and renewal model inference problems. The left panels (a) show how the reconstructed phylogeny of infecteds (i) leads to (branching) coalescent events, which form the Poisson count record of (ii). The timing of these observable events encodes information about the piecewise effective population size function to be inferred in (iii). The right panels (b) indicate how infecteds, which naturally conform to the Poisson count record of (i) are usually only observed at the resolution of days or weeks, leading to the Poisson histogram record in (ii). The number of infecteds in these histogram bins inform on the piecewise effective reproduction number in (iii). Both models feature data with *m* = 4 and involve *p* = 2 parameters to be estimated. See Materials and Methods for notation.

The effective reproduction number at time *s*, *R*(*s*), is a key diagnostic of whether an outbreak is growing or under control. It defines how many secondary infections an infected will, on average, generate [43]. The renewal or branching process model [7] is a popular approach for inferring *R*(*s*) from epicurves that generalises the Lotka-Euler equation from ecology [42]. Renewal models describe how fluctuations in *R*(*s*) modulate the tree-like propagation structure of an epidemic and have been used to predict Ebola virus disease case counts and assess the transmissibility of pandemic influenza, for example [8] [4] [23]. Here *s* indicates discrete time e.g. days.

The effective population size at *t*, *N*(*t*), is a popular proxy for census (or true) population size that derives from the genetic diversity of the target demography. When applied to epidemics, *N*(*t*) measures the number of infecteds contributing offspring (i.e. transmitting the disease) to the next generation [13]. The skyline plot model [32] is a prominent means of estimating *N*(*t*) from phylogenies that extends the Kingman coalescent process from population genetics [16]. Skyline models explain how variations in *N*(*t*) influence the shape and size of the infected genealogy and have informed on the historical transmission and origin of HIV, influenza and hepatitis C, among others [31] [18] [34]. Here *t* is continuous and usually in units of genealogical time.

While renewal and skyline models depict very different aspects of an infectious disease, they possess some statistical similarities. Foremost is their approximation of *N*(*t*) and *R*(*s*) by *p*-dimensional, piecewise-constant functions (see (iii) in Fig. 1). Here *p* is the number of parameters to be inferred from the data and time is regressive for phylogenies but progressive for incidence curves. The choice of *p* is critical to the quality of inference. Models with large *p* can better track rapid changes but are susceptible to noise and uncertainty (overfitting) [4]. Smaller *p* improves estimate precision but reduces flexibility, easily over-smoothing (underfitting) salient changes [20]. Optimally selecting *p*, in a manner that is justified by the available data, is integral to deriving reliable and sensible conclusions from these models.

Surprisingly, no transparent, principled and easily computable *p*-selection strategy exists. In renewal models, *p* is often set by trial and error, or defined using heuristic sliding windows [4] [7]. Existing theory on window choice is limited, with [4] positing a bound on the minimum number of infecteds a window should contain for a given level of estimate uncertainty and [23] initially proposing a ‘naïve-rational’ squared error based window-sizing approach, which they subsequently found inferior to other subjective window choices examined in that study. In skyline models, this problem has been more actively researched because the classic skyline plot [32], which forms the core of most modern skyline methods, overfits by construction i.e. it infers a parameter per data-point. Accordingly, various approaches for reducing *p*, by ensuring that each population size parameter is informed by groups of data-points, have been proposed.

The generalised skyline plot [38] uses a small sample correction to the Akaike information criterion (AIC) to achieve one such grouping in an interpretable and computable fashion. However, basing analyses solely on the AIC can still lead to overfitting [15]. The Bayesian skyline plot built on the generalised skyline by additionally incorporating a prior distribution that assumed an exponentially distributed autocorrelation between successive parameters [6]. This implicitly influenced group choices but is known to oversmooth or underfit [20]. As a result, later approaches such as the Skyride and Skygrid reverted to the classic skyline plot and applied Gaussian-Markov smoothing prior distributions to achieve implicit grouping [20] [9]. However, these methods also raised concerns about underfitting and the relationship between model selection and smoothing prior settings is obscure [29].

Other approaches to effective population size model selection are considerably more involved. The extended Bayesian skyline plot and multiple change-point method use piecewise-linear functions and apply Bayesian stochastic search variable selection [12] and reversible jump MCMC [24] to optimise *p*. These algorithms, while capable, are more computationally demanding, and lack interpretability (their results are not easily debugged and linear functions do not possess the biological meaningfulness of constant ones, which estimate the harmonic mean of time-varying population sizes [32]). Note that we assume phylogenetic data is available without error (i.e. we do not consider extensions of the above or subsequent methods to genealogical uncertainty) and limit the definition of skyline models to those with piecewise-constant functions. In Fig. A4 of the Appendix we illustrate estimates from some of these approaches on an HIV dataset.

New *p*-selection metrics, which can balance between the interpretability of the generalised skyline and the power of more sophisticated Bayesian selection methods, are therefore needed. Here we attempt to answer this need by developing and validating a minimum description length (MDL)-based approach that unifies renewal and skyline model selection. MDL is a formalism from information theory that treats model selection as equivalent to finding the best way of compressing observed data (i.e. its shortest description) [35]. MDL is advantageous because it includes both model dimensionality and parametric complexity within its definition of model complexity [36]. Parametric complexity describes how the functional relationship between parameters matters [21], and is usually ignored by standard selection criteria. However, MDL is generally difficult to compute [10], which may explain why it has not penetrated the epidemiological or phylodynamics literature.

We overcome this issue by deriving a tractable Fisher information approximation (FIA) to MDL. This is achieved by recognising that sampled phylogenies and incidence curves both sit within a Poisson point process framework and by capitalising on the piecewise-constant structure of skyline and renewal models. The result is a pair of analogous FIA metrics that lead to adaptive estimates of *N*(*t*) and *R*(*s*) by selecting the *p* most justified by the observed Poisson data. These expressions decompose model complexity into clearly interpretable contributions and are as computable as the standard AIC and the Bayesian information criteria (BIC). We find, over a range of selection problems, that the FIA generally outperforms the AIC and BIC, emphasising the importance of including parametric complexity. This improvement requires some knowledge about the piecewise parameter space domain.

## II. Materials and Methods

### A. Phylogenetic Skyline and Epidemic Renewal Models

The phylogenetic skyline and epidemic renewal models are popular approaches to solving epidemiological inference problems. The skyline plot or model [13] infers the hidden, time-varying effective population size, *N*(*t*), from a phylogeny of sequences sampled from that infected population; while the renewal or branching process model [8] estimates the hidden, time-varying effective reproduction number, *R*(*s*), from the observed incidence of an infectious disease. Here *t* indicates continuous time, which is progressive (moving from past to present) in the renewal model, but reversed (retrospective) in the skyline. We initially use *R*(*t*) here before deriving *R*(*s*). While both models solve different problems, they approximate their variable of interest, *θ*(*t*), with a *p*-dimensional piecewise-constant function, and assume a Poisson point process relationship between it and the observed data, *Y*(*t*), as in Eq. (1).

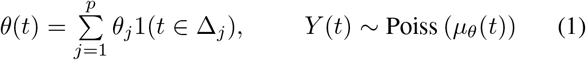

Here *θ*(*t*) is either *N*(*t*) or *R*(*t*) and *Y*(*t*) is either phylogenetic or incidence data, depending on the model of interest. The *j*^th^ piecewise component of *θ*(*t*), which is valid over the closed interval Δ_*j*_, is *θ_j_*. The rate function, *μ_θ_*(*t*) depends on *θ*(*t*) and allows us to treat the usually distinct skyline and renewal models within the same Poisson point process framework. We want to estimate the parameter vector *θ* = [*θ*_1_, …, *θ_p_*] from the data over 0 ≤ *t* ≤ *T*, denoted 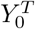. We consider two fundamental mechanisms for observing 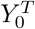 and then show how they apply to skyline and renewal models.

The first, known as a Poisson count record [37], involves having access to every event time of the Poisson process i.e. 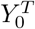 is observed directly. Eq. (2) gives the likelihood of these data, in which a total of *m* events occur.

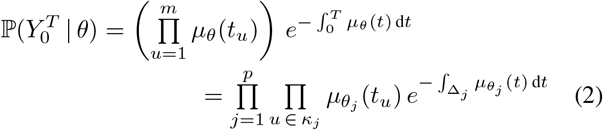

The *u*^th^ event time is *t_u_* and 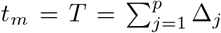. The set *κ_j_* = {*u* : *t_u_* ∈ Δ_*j*_} collects all event indices within the *j*^th^ piecewise interval and 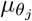 emphasises that the parameter controlling the rate in Δ_*j*_ is *θ_j_*. We denote the portion of events falling within Δ_*j*_ as *m_j_* so that 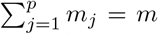. The number of elements in *κ_j_* is therefore *m_j_* with the first and last of these *m_j_* event times delimiting Δ_*j*_. The size of the data is also summarised by *m*.

The second is called a Poisson histogram record [37] and applies when individual events are not observed. Instead only counts of the events occurring within time bins are available and the size of the data is now defined by the number of bins. We redefine *m* for this data type as the number of bins so that it again controls data size. The *s*^th^ bin has width *b_s_* and count *c_s_* and we use 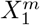 to denote the bin transformed version of 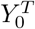. The likelihood is then given by Eq. (3).

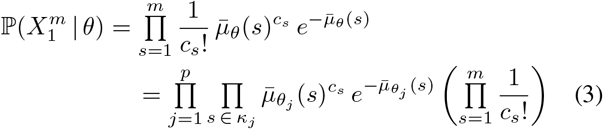

Here 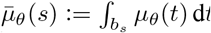 is the Poisson rate integrated across the *s*^th^ observation bin and *κ_j_* again defines the indices (of bins in this case) that are controlled by *θ_j_*. The time interval over which *θ_j_* is valid is 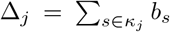. Fig. 1 illustrates the relationship between histogram and count records. We now detail how these observation schemes apply to phylogenetic and incidence data and hence skyline and renewal models.

The skyline model is based on the coalescent approach to phylogenetics [16]. Here genetic sequences (lineages) sampled from an infected population across time elicit a reconstructed phylogeny or tree, in which these lineages successively merge into their common ancestor. The observed branching or coalescent times of this tree form a Poisson point process that contains information about the piecewise effective population parameters *N* ≔ [*N*_1_*, …, N_p_*]. Since the coalescent events times, {*t_u_*} are observable, phylogentic data correspond to a Poisson count record. The rate underlying the events for *t* ∈ Δ_*j*_ is 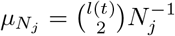 with *l*(*t*) counting the lineages in the phylogeny at time *t* (this increases at sample event times and decrements at coalescent times).

The log-likelihood of the observed, serially sampled tree data, denoted by count record 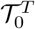 is then derived from Eq. (2) to obtain Eq. (4), which is equivalent to standard skyline log-likelihoods [6], but with constant terms removed.

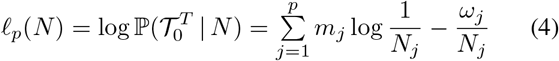

Here 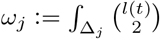 d*t* and *m_j_* counts the number of coalescent events falling within Δ_*j*_. The endpoints of Δ_*j*_ coincide with coalescent event times, as in [32] [6] [25]. Fig. 1a outlines the skyline coalescent inference problem and summarises its notation. Since *N*(*t*) can have a large dynamic range (e.g. for exponentially growing epidemics) we will analyse the skyline model under the robust log transform [28], which ensures good statistical properties.

The maximum likelihood estimate (MLE), and Fisher information (FI) are important measures for describing how estimates of *N* (or log *N*) depend on 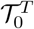. We compute the MLE, 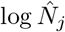, and FI, 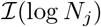, of the skyline model by solving 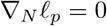 and 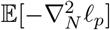 and then log-transforming, with ∇_*N*_ ≔ {*∂/∂N*_*j*_} as the vector derivative operator [17]. The result is Eq. (5) [28].

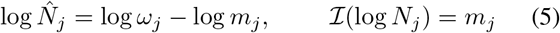

For a given *p*, the MLE controls the per-segment bias because as *m*_*j*_ increases 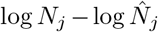 decreases. The FI defines the precision i.e. the inverse of the variance around the MLEs, and also (directly) improves with *m_j_*. We will find these two quantities to be integral to formulating our approach to *p*-model selection. The FI and MLE control the per-segment performance, while *p* determines how well the overall piece-wise function adapts to the underlying generating process.

The renewal model is based on the classic (Lotka-Euler) renewal equation or branching process approach to epidemic transmission [42]. This states that the number of new infecteds depends on past incidence through the generation time distribution, and the effective reproduction number *R*(*s*). As incidence is usually observed on a coarse temporal scale (e.g. days or weeks), exact infection times are not available. As a result, incidence data conform to a Poisson histogram record with the number of infecteds observed in the *s*^th^ bin denoted *I*(*s*). For simplicity we assume daily bins. The generation time distribution is specified by *w*(*u*), the probability that an infected takes between *u* − 1 and *u* days to transmit that infection [7].

The total infectiousness of the disease is 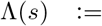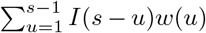. We make the common assumptions that *w*(*u*) is known (it is disease specific) and stationary (does not change with time) [4]. If an epidemic is observed for *m* days then the historical incidence counts, 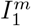, constitute the histogram record informing on the piecewise parameters to be estimated, *R* = [*R*_1_, …, *R_p_*]. The renewal equation asserts that 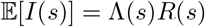 [7]. Setting this as the integrated bin rate 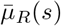 leads to the log-likelihood of Eq. (6) from Eq. (3).

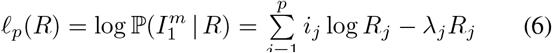

Here 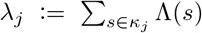 and 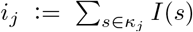 are sums across the indices *κ_j_*, which define the *m_j_* bins composing Δ_*j*_. Eq. (6) is equivalent to the standard renewal log-likelihood [8] but with the constant terms removed.

This derivation emphasises the statistical similarity between count and histogram records (and hence skyline and renewal models) and allows generalisation to variable width histogram records (e.g. irregularly timed epi-curves). Fig. 1b illustrates the renewal inference problem and its notation. We can compute the MLE and robust FI from Eq. (6) as Eq. (7) [8] [28].

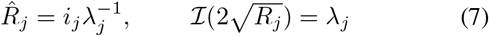

As each *m_j_* becomes large the per-segment bias 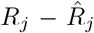 decreases. Using results from [28], we find the square root transform of *R* to be robust for renewal models i.e. it guarantees optimal estimation properties. We compute the FI under this parametrisation to reveal that the total infectiousness controls the precision around our MLEs (via *λ_j_*). This will also improve as *m_j_* increases, but with the caveat that the parameters underlying bigger epidemics (specified by larger historical incidence values, and controlled through Λ(*s*)) are easier to estimate than those of smaller ones.

In both models we therefore find a clear piecewise separation of MLEs and FIs. Per-segment bias and precision depend on the quantity of data apportioned to each parameter. This data division is controlled by *p*, which balances per-segment performance against the overall fit of the model to its generating process. Thus, model dimensionality fundamentally controls inference quality. Large *p* means more segments, which can adapt to rapid *N*(*t*) or *R*(*s*) changes. However, this also rarefies the per-segment data (grouped sums like *λ_j_* or *m_j_* decrease) with both models becoming unidentifiable if *p > m*. Small *p* improves segment inference, but stiffens the model. We next explore information theoretic *p*-selection methods that formally utilise these MLEs and FIs in their decision making.

#### Model and Parametric Complexity

Our proposed approach to model selection relies on the MDL framework of [35]. This treats modelling as an attempt to compress the regularities in the observed data, which is equivalent to learning about its statistical structure. MDL evaluates a *p*-parameter model, 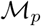, in terms of its code length (in e.g. nats or bits) as 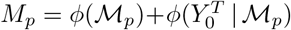 [10]. Here *ϕ*(*x*) computes the length to encode *x* and 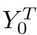 is the observed data. *M_p_* is the sum of the information required to describe 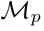 and the data given that 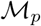 is chosen. More complex models have larger 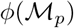 (more bits are needed to describe the model), and smaller 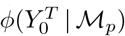 (as complex models can better fit the data, there is less remaining information to detail).

If *n* models are used to describe 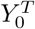 then the model with *p*\* = arg min_1≤*p*≤*n*_ *M*_*p*_ best compresses or most succinctly represents the data. The model with *p*\* is known to possess the desirable properties of generalisability and consistency [10]. The first means that 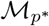 provides good predictions on newly observed data (i.e. it fits the underlying data generating process instead of a specific instance of data from that process), while the second indicates that the selected *p*\* will converge to the true model index (if one exists) as data increase [30] [1]. If *θ* represents the *p*-parameter vector of 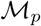 and 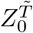 is a potential instance of data derived from the same generating process as *Y* then the MDL code lengths can be reframed as 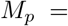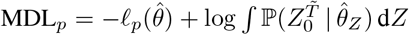 [36].

The first term of MDL_*p*_ describes the goodness-of-fit of the model to the observed data, while the second term balances this against the fit to unobserved data (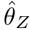 is the MLE of the parameters of 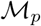 but with 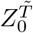 as data) from the same process. This is done over all possible data that could be obtained from that process (hence the integral with respect to d*Z*) and measures the generalisability of the model. This generalisability term is usually intractable. We therefore use a well known FI approximation from [36], which we denote FIA*_p_* 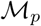 in Eq. (8), with ‘det’ as the matrix determinant.

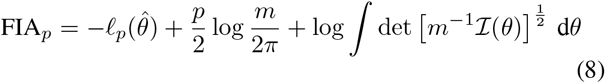

The approximation of Eq. (8) is good, provided certain regularity conditions are met. These mostly relate to the FI being identifiable and continuous in *θ*, and are not issues for either skyline or renewal models [21]. While we will apply the FIA within a class of renewal or skyline models, this restriction is unnecessary. The FIA can be used to select among any variously parametrised and non-nested models [10].

The FIA not only maintains the advantages of MDL, but also has strong links to Bayesian model selection (BMS). BMS compares models based on their posterior evidence i.e. 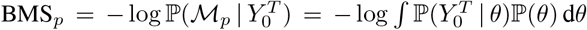 [15]. BMS and MDL are considered the two most complete and rigorous model selection measures [10]. As with MDL, the BMS integral is often intractable and it can be difficult to disentangle and interpret how the formulation of 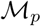 impacts its associated complexity according to these metrics [30]. Interestingly, if a Jeffreys prior distribution is used for 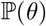, then it can be shown that BMS_*p*_ ≈ FIA_*p*_ (via an asymptotic expansion) [21]. Consequently, the FIA uniquely trades off between the performance of BMS and MDL for some computational ease.

However, this trade-off is not perfect. For many model classes the integral of the FI in Eq. (8) can be divergent or difficult to compute [10]. At the other end of the computability–completeness spectrum are standard metrics such as the AIC and BIC, which are quick and simple to construct, calculate and interpret. These generally penalise a goodness-of-fit term (e.g. 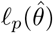) with the number of parameters *p* and may also consider the total size of the data *m*. Unfortunately, these methods often ignore the parametric complexity of a model, which measures the contribution of the functional form of a model to its overall complexity. Parametric complexity explains why two-parameter sinusoidal and exponential models have non-identical complexities, for example. This concept is detailed in [30] and [10] and corresponds to the FI integral.

This provides the statistical context for our proposing the FIA as a meaningful metric for skyline and renewal models. In Results we will show that the piecewise separable MLEs and FIs (Eq. (5) and Eq. (7)) of these models not only ensure that the FI integral is tractable, but also guarantee that Eq. (8) is no more difficult to compute than the AIC or BIC. Consequently, our proposed adaptation of the FIA is able to combine the simplicity of standard measures such as the AIC and BIC while still capturing the more sophisticated and comprehensive descriptions of complexity inherent to the BMS and MDL by including parametric complexity. This point is embodied by the relationship between the FIA and BIC. As data size asymptotically increases the parametric complexity becomes less important (it does not grow with *m*) and FIA_*p*_ → BIC_*p*_. The BIC is hence a coarser approximation to both the MDL and BMS, than the FIA [21].

While the FIA achieves a favourable compromise among interpretability, completeness and computability in its description of complexity, it does depend on roughly specifying the domain of the FI integral. We will generally assume some arbitrary but sensible domain. However, when this is not possible the Qian-Kunsch approximation to MDL, denoted QK_*p*_ and given in Eq. (9), can be used [33].

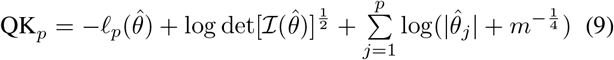

This approximation trades off some interpretability and performance for the benefit of not having to demarcate the multidimensional domain of integration.

Lastly, we provide some intuition about Eq. (8), which balances fit via the maximum log-likelihood 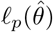 against model complexity, which can be thought of as a geometric volume defining the set of distinguishable behaviours (i.e. parameter distributions) that can be generated from the model. This volume is composed of two terms. The first, 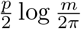, shows, unsurprisingly, that higher model dimensionality, *p*, expands the volume of possible behaviours. Less obvious is the fact that increased data size *m* also enlarges this volume because distinguishability improves with inference resolution. The second term, which is parametric complexity, is invariant to transformations of *θ*, independent of *m* and is an explicit volume integral measuring how different functional relationships among the parameters, defined via the FI, influence the possible, distinguishable behaviours the model can depict [10].

## III. Results

### The Insufficiency of Log-Likelihoods

The inference performance of both the renewal and skyline models, for a given dataset, strongly depends on the chosen model dimensionality, *p*. As observed previously, current approaches to *p*-selection utilise ad-hoc rules or elaborate algorithms that are difficult to interrogate. Here we emphasise why finding an optimal *p*, denoted *p*\*, is important and illustrate the pitfalls of inadequately balancing bias and precision. We start by proving that overfitting is a guaranteed consequence of depending solely on the log-likelihood for *p*-selection. While this may seem obvious, early formulations of piecewise models did over-parametrise by setting *p* = *m* [38] and our proof can be applied more generally e.g. when selecting among models with *p* < *m*. Substituting the MLEs of Eq. (5) and Eq. (7) into Eq. (4) and Eq. (6), we get Eq. (10).

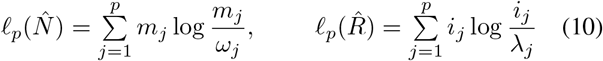

Both the renewal and skyline log-likelihoods take the form 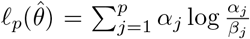, due to their inherent and dominant piecewise-Poisson structure. Here *α*_*j*_ and *β*_*j*_ are grouped variables that are directly computed from the observed data (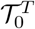 or 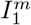). The most complex model supportable by the data is at *p* = *m* with 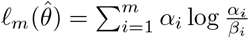 As the date size (*m*) is fixed we can clump the *i* indices falling within the duration of the *j*^th^ group Δ_*j*_ as 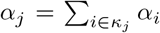 and 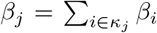. The log-sum inequality from [5] status that 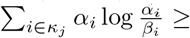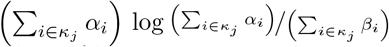. Repeating this across all possible *p* groupings results in Eq. (11).

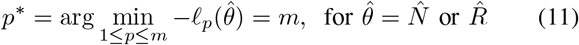

Thus, log-likelihood based model selection always chooses the highest dimensional renewal or skyline model. This result also holds when solving Eq. (11) over a subset of all possible *p*, provided smaller *p* models are non-overlapping groupings of larger *p* ones [11]. Thus, it is necessary to penalise 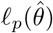 with some term that increases with *p*.

The highest *p*-model is most sensitive to changes in *θ*(*t*), but extremely noisy and likely to overfit the data. This noise is reflected in a poor FI. From Eq. (5) and Eq. (7) it is clear that grouping linearly increases the FI, hence smoothing noise. However, this improved precision comes with lower flexibility. At the extreme of *p* = 1, for example, *θ*(*t*) is approximated by a single, perennial parameter, and the log-likelihood 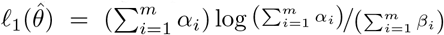 is unchanged for all combinations of data that produce the same grouped sums. This oversmooths and underfits. We will always select *p*\* = 1 if our log-likelihood penalty is too sensitive to dimensionality.

We now present some concrete examples of bad model selection. We use adjacent groupings of size *k* to control *p* i.e. every *κ*_*j*_ clumps *k* successive indices (the last index is *m*). In Fig. 2a we examine skyline models with periodic exponential fluctuations ((i)-(ii)) and bottleneck variations ((iii)-(iv)). The periodic case describes seasonal epidemic oscillations in in fecteds, while the bottleneck simulates the severe decline that results from a catastrophic event. In Fig. 2b we investigate renewal models featuring cyclical ((i)-(ii)) and sigmoidal ((iii)-(iv)) *R*(*s*) dynamics. The cyclical model depicts the pattern of spread for a seasonal epidemic (e.g. influenza), while the sigmoidal one portrays a vaccination policy that quickly leads to outbreak control.

**Fig. 2:**
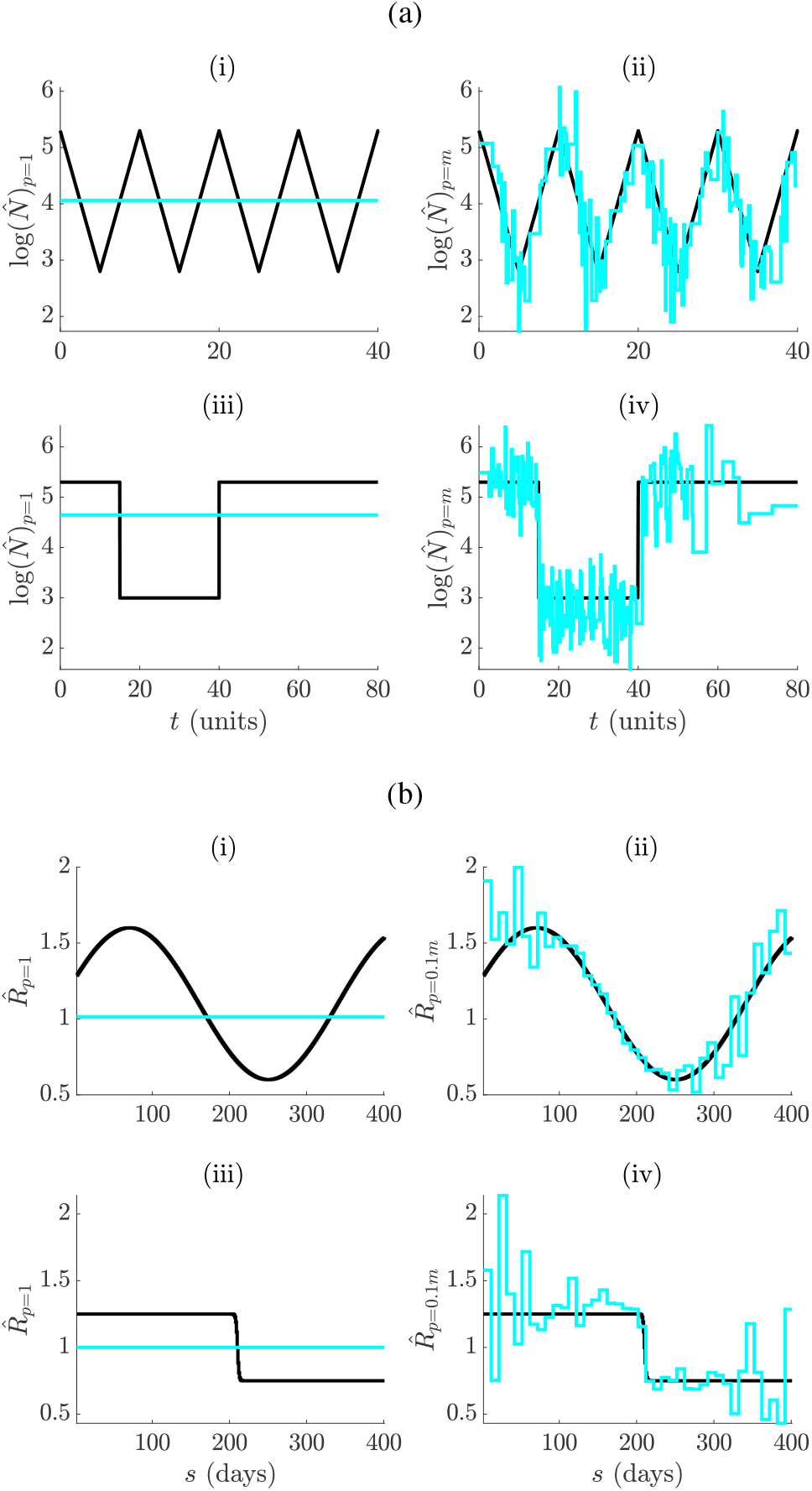
Skyline and renewal model under and overfitting. Small *p* leads to smooth but biased estimates characteristic of underfitting ((i) and (iii) in (a) and (b)). Large *p* results in noisy estimates that respond well to changes. This is symptomatic of overfitting ((ii) and (iv) in (a) and (b)). The MLEs (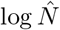 or 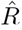) are in cyan and the true log *N*(*t*) or *R*(*s*) in black. Panel (a) shows cyclic and bottleneck skyline models at *m* = 800 and (b) focuses on sinusoidal and sigmoidal renewal models at *m* = 400.

In both Fig. 2a and Fig. 2b we observe underfitting at low *p* ((i) and (iii)) and overfitting at high *p* ((ii) and (iv)). The detrimental effects of choosing the wrong model are not only dramatic, but also realistic. For example, in the skyline examples the underfitted case corresponds to the fundamental Kingman coalescent model [16], which is often used as a null model in phylogenetics. Alternatively, the classic skyline [32], which is at the core of many coalescent inference algorithms, is exactly as noisy as the overfitted case. Correctly penalising the log-likelihood is therefore essential for good estimation, and forms the subject of the subsequent section.

### Minimum Description Length Selection

Having clarified the impact of non-adaptive estimation, we develop and appraise various, easily-computed, model selection metrics, in terms of how they penalise renewal and skyline log-likelihoods. The most common and popular metrics are the AIC and BIC [15], which we reformulate in Eq. (12) and Eq. (13), with (*α*_*j*_, *β*_*j*_) = (*m*_*j*_, *ω*_*j*_) or (*i*_*j*_, *λ*_*j*_) for skyline and renewal models respectively.

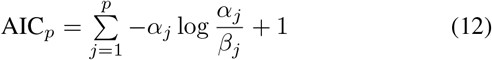

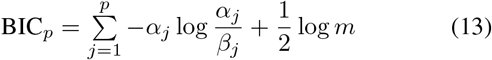

By decomposing the AIC and BIC on a per-segment basis (for a model with *p* segments or dimensions), as in Eq. (12) and Eq. (13), we gain insight into exactly how they penalise the log-likelihood. Specifically, the AIC simply treats model dimensionality as a proxy for complexity, while the BIC also factors in the total dimension of the available data. A small sample correction to the AIC, which adds a further *p*+1/*m*−*p*−1 to the penalty in Eq. (12), was used in [38] for skyline models. We found this correction inconsequential to our later simulations and so used the standard AIC only.

As discussed in Materials and Methods, these metrics are insufficient descriptions because they ignore parametric complexity. Consequently, we suggested the MDL approximations of Eq. (8) and Eq. (9). We now derive and specialise these expressions to skyline and renewal models. Adapting the FIA metric of Eq. (8) forms a key result of this work. Its integral term, 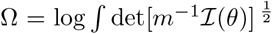, can, in general, be intractable [36]. However, the piecewise structure of both the skyline and renewal models, which leads to orthogonal (diagonal) FI matrices, allows us to decompose det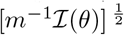 as 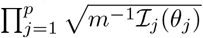 with 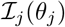 as the *j*^th^ diagonal element of 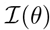, which only depends on *θ*. Note that *θ* = N or R for the skyline and renewal model respectively.

Using this decomposition we can partition Ω across each piecewise segment as 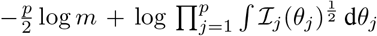. The 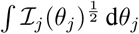 is known to be invariant to parameter transformations [10]. This is easily verified by using the FI change of variable formula [17]. This asserts that 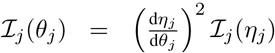, with *η*_*j*_ as some function of *θ*_*j*_. The orthogonality of our piecewise-constant FI matrices allows this component-by-component transformation. Hence 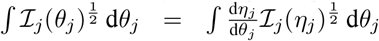, which equals 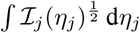. We let *η*_*j*_ denote the robust transform of 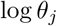 or 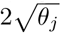 for the skyline or renewal model, respectively. Robust transforms make the integral more transparent by removing the dependence of 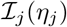 on *η_j_* [28].

Hence we use Eq. (5) 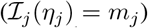 and Eq. (7) 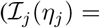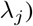 to further obtain 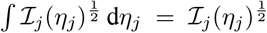 and 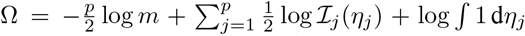. The do main of integration for each parameter is all that remains to be solved. We make the reasonable assumption that each piecewise parameter, *θ*_*j*_, has an identical domain. This is *N*_*j*_ [1, *ν*] and *R*_*j*_ ∈ [0, *ν*], with *ν* as an unknown model-dependent maximum. The minima of 1 and 0 are sensible for these models. This gives 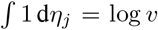 or 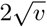 for the skyline or renewal model. Substituting into Ω and Eq. (8) yields Eq. (14) and Eq. (15).

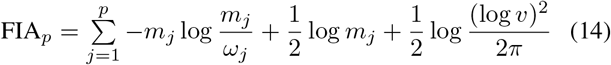

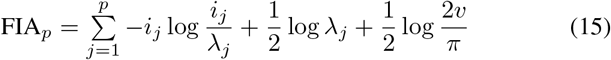

Eq. (14) and Eq. (15) present an interesting and more complete view of piecewise model complexity. Comparing to Eq. (13) reveals that the FIA further accounts for how the data are divided among segments, making explicit use of the robust FI of each model. This is an improvement over simply using the (clumped) data dimension *m*. Intriguingly, the maximum value of each parameter to be inferred, *ν*, is also central to computing model complexity. This makes sense as models with larger parameter spaces can describe more types of dynamical behaviours [10]. By comparing these terms we can disentangle the relative contribution of the data and parameter spaces to complexity.

One limitation of the FIA is its dependence on the unknown *ν*, which is assumed finite. This is reasonable as similar assumptions would be implicitly made to compute the BMS or MDL (in cases where they are tractable). The QK metric [33], which also approximates the MDL, partially resolves this issue. We compute QK_*p*_ by substituting FIs and MLEs into Eq. (9). Expressions identical to Eq. (14) and Eq. (15) result, except for the *v*-based terms, which are replaced as in Eq. (16) and Eq. (17).

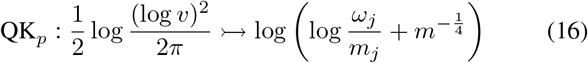

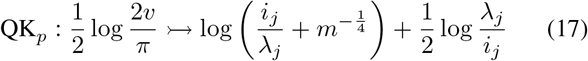

These replacements require no knowledge of the parameter domain, but still approximate the parametric complexity of the model [33]. However, in gaining this domain independence we lose some performance (see later sections), and transparency. Importantly, both the FIA and QK are as easy to compute as the AIC or BIC. The similarity in the skyline and renewal model expressions reflects the significance of their piecewise-Poisson structure. We next investigate the practical performance of these metrics.

### Adaptive Estimation: Epidemic Renewal Models

We validate our FIA approach on several renewal inference problems. We simulate incidence curves, 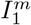, via the renewal or branching process relation *I*(*s*) ~ Poiss(Λ(*s*)*R*(*s*)) with *R*(*s*) as the true effective reproduction number that we wish to estimate. We construct Λ(*s*) using a gamma generation time distribution that approximates the one used in [23] for Ebola virus outbreaks. We initialise each epidemic with 10 infecteds as in [4]. We condition on the epidemic not dying out, and remove initial sequences of zero incidence to ensure model identifiability. We consider an observation period of *m* = 400 days, and select among models with 10 ≤ *k* ≤ *m* m such that *m* is divisible by *k*. Here *k* counts how many days are grouped to form a piecewise segment (i.e. the size of every *κ*_*j*_), and model dimensionality, *p*, is bijective in *k* i.e. *pk* = *m*.

We apply the criteria developed above to select among possible *p*-parameter (or *k*-grouped) renewal models. For the FIA we set *ν* = 100 as a conservative upper bound on the reproduction number domain. We start by highlighting how the FIA (1) regulates between the over and underfitting extremes from Fig. 2b, and (2) updates its selected *p*\* as the data increase. These points are illustrated in Fig. 3a and Fig. 3b. Graphs (i) and (iii) exemplify (1) as the FIA ((iii)) reduces *p* from the maximum chosen by the log-likelihood ((i)), leading to estimates that balance noise against dimensionality. Interestingly, the FIA chooses a minimum of segments for the sigmoidal fall in Fig. 3b, and so pinpoints its key dynamics. As the observed data are increased (graphs (ii) and (iv) of Fig. 3a and Fig. 3b) the FIA adapts *p* to reflect the improved resolution that is now justified, hence demonstrating (2). The increased data use 5 more, conditionally independent (on *R(s))* 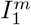 curves and has size 6*m*. The *i*_*j*_ and *λ*_*j*_ used now sum over every of the 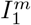 curves.

**Fig. 3:**
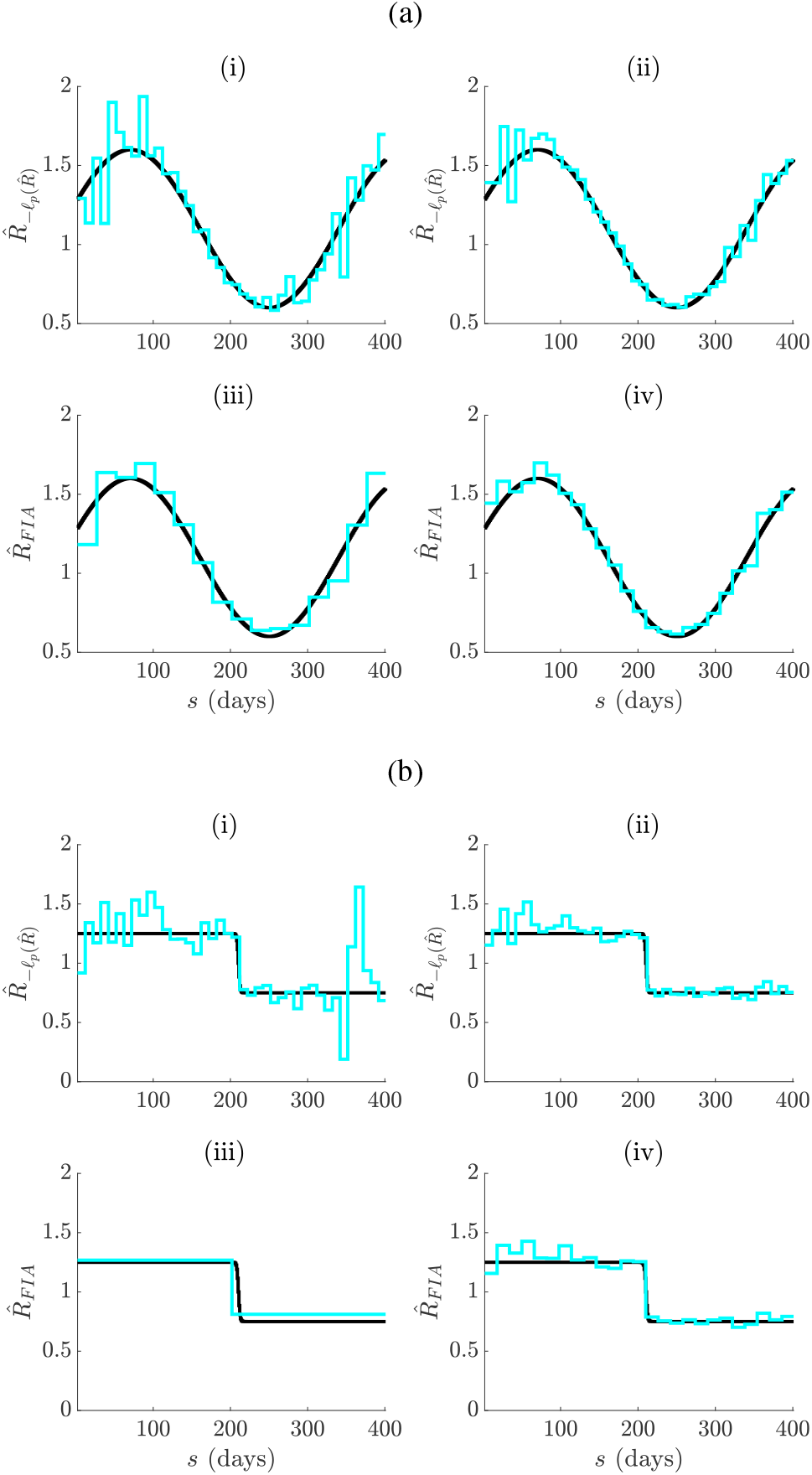
Adaptive cyclical and sigmoidal estimation with FIA. In (a) and (b), graphs (i)-(ii) present optimal log-likelihood based *R*(*s*) MLEs for 1 ((i)) and 6 ((ii)) observed incidence data streams, simulated under renewal models with time-varying effective reproduction numbers. Graphs (iii)-(iv) give the FIA adaptive estimates at the same settings with *ν* = 100. Panels (a) and (b) examine cyclical and sigmoidal (also called logistic) reproduction number profiles, respectively.

While the above examples provide practical insight into the merits of the FIA, they cannot rigorously assess its performance, since continuous *R*(*s*) functions have no true *p* = *p*\* or *k*\* = *m/p*\*. We therefore study two problems in which a true *p*\* exists: a simple binary classification, and a more complex piecewise model search. In both, we benchmark the FIA against the AIC, BIC and QK criteria over the same set of simulated 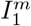 curves. We note that, when *R*(*s*) is piecewise-constant, increasing the number of conditionally independent curves improves the probability of recovering *p*\*. We discuss the results of the first problem in the Appendix (see Fig. A1), where we show that the FIA most accurately identifies between a null model of an uncontrolled epidemic and an alternative model featuring rapid outbreak control. The FIA uniformly outperforms all other metrics at every *p*\* in this problem, with the QK a close second.

For the second, more complicated problem, we consider models involving piecewise-constant *R*(*s*) changes after every *k*\* days, with *k*\* looping over 20 ≤ *k* ≤ *m* and *p*\**k*\* = *m* = 400 days. For every *k*\* we generate 10^5^ independent epidemics, allowing *R*(*s*) to vary in each run, with magnitudes uniformly drawn from [*R*_min_, *R*_max_]. Fig. 4a illustrates typical random telegraph *R*(*s*) models at each *k*\* (these change in magnitude for each run). Key selection results are shown in Fig. 4b with *R*_min_ = 0.5, (*R*_max_, *ν*) = [5, 100] in (i) and [1.5, 1.5] in (ii). In both cases, the FIA attains the best overall accuracy i.e. 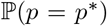, followed by the QK (which overlaps the FIA curve in (i)), BIC and AIC. The dominance of both MDL-based criteria suggests that parametric complexity is important. However, the FIA can do worse than the BIC and QK when *ν* is large compared to *R*_max_ (or if Rmin is notably above 0). We discuss these cases in the Appendix (see Fig. A3), explaining why *ν* = 1:5 is used in (ii).

**Fig. 4:**
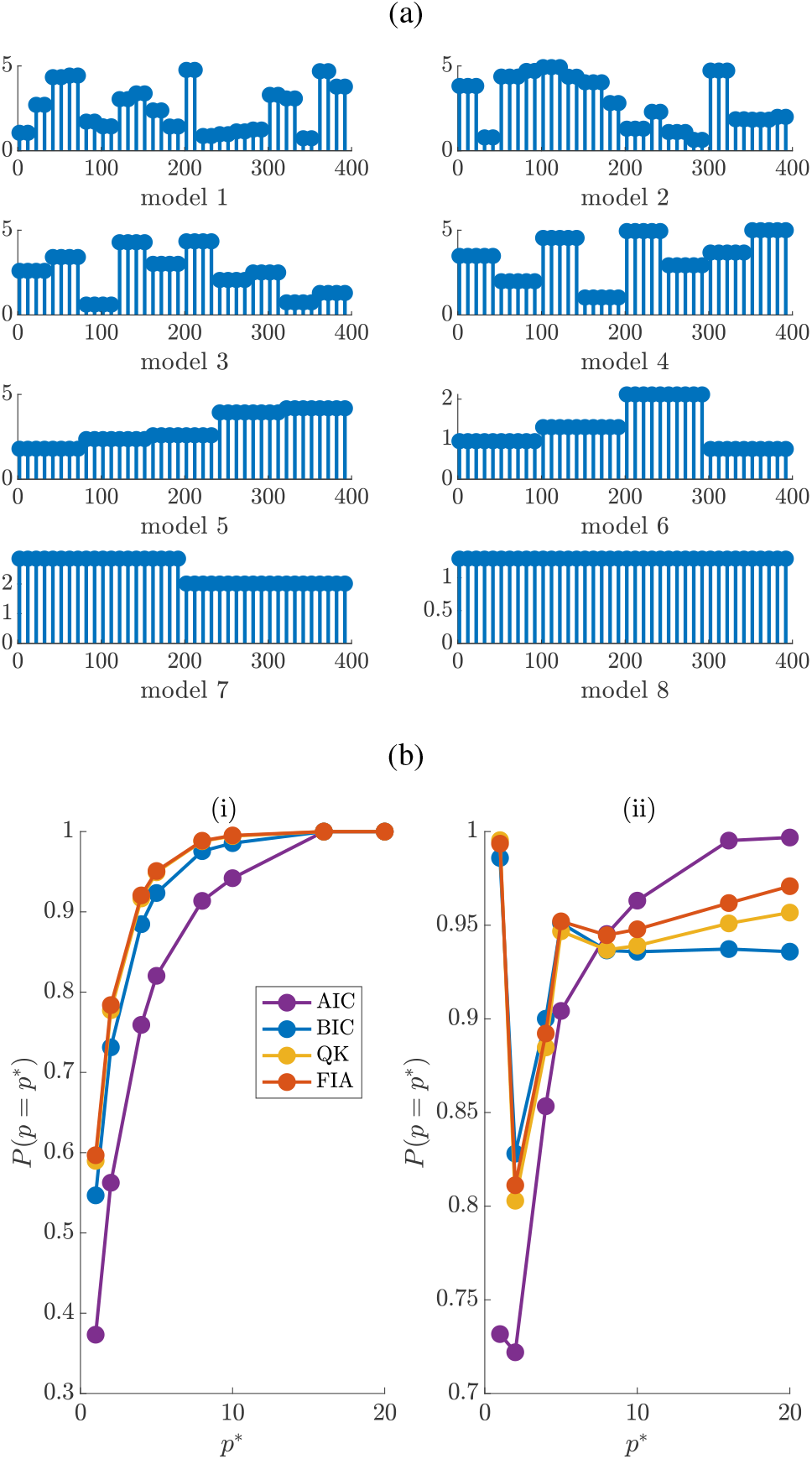
Renewal model selection. We simulate 10^5^ epidemics from renewal models with 20 ≤ *k* ≤ *m* = 400 and *p* = *m/k*. We test the ability of several model selection criteria to recover the true *p* = *p*\* from this set. Each epidemic has an independent, piecewise-constant *R*(*s*), examples of which are shown in (a). These models change in amplitude but not *k* for every simulation. Panel (b) shows the probability of detecting the true model against *p*\* and (i) considers *R*(*s*) ∈ [0.5, 5] with *ν* = 100 while (ii) uses *R*(*s*) ∈ [0.5, 1.5] and *ν* = 1.5. The FIA performs best at every *p*\* in (i) and overall in (ii).

### Adaptive Estimation: Phylogenetic Skyline Models

We verify the FIA performance on several skyline problems. We simulate serially sampled phylogenies with sampled tips spread evenly over some interval using the phylodyn R package of [14]. Increasing the sampling density within that interval increases overall data size *m* (each pair of sampled tips can produce a coalescent event). We define our *p* segments as groups of *k* coalescent events. Skyline model selection is more involved because the end-points of the *p* segments coincide with coalescent events. While this ensures statistical identifiability, it means that grouping is sensitive to phylogenetic noise [38], and that *p* changes for a given *k* if *m* varies (*m* = *pk*). This can result in MLEs, even at optimal groupings, appearing delayed or biased relative to *N*(*t*), when *N*(*t*) is not a grouped piecewise function. Methods are currently under development to resolve these biases [25].

Nevertheless, we start by examining how our FIA approach mediates the extremes of Fig. 2a. We restrict our grouping parameter to 4 ≤ *k* ≤ 80, set *ν* = 10^3^ (max *N*(*t*) = 300) and apply the FIA of Eq. (14) to obtain Fig. 5a and Fig. 5b. Two points are immediately visible: (1) the FIA ((iii)-(iv)) regulates the noise from the log-likelihood ((i)-(ii)), and (2) the FIA supports higher *p*\* when the data are increased ((iv)). Specifically, the FIA characterises the bottleneck of Fig. 5b using a minimum of segments but with a delay. As data accumulate, more groups can be justified and so the FIA is able to compensate for the delay. Note that the last 1-2 coalescent events are often truncated, as they can span half the time-scale, and bias all model selection criteria [22]. In the Appendix (see Fig. A4) we show how the sensitivity of the FIA to event density compares to other methods on empirical data (see Materials and Methods).

**Fig. 5:**
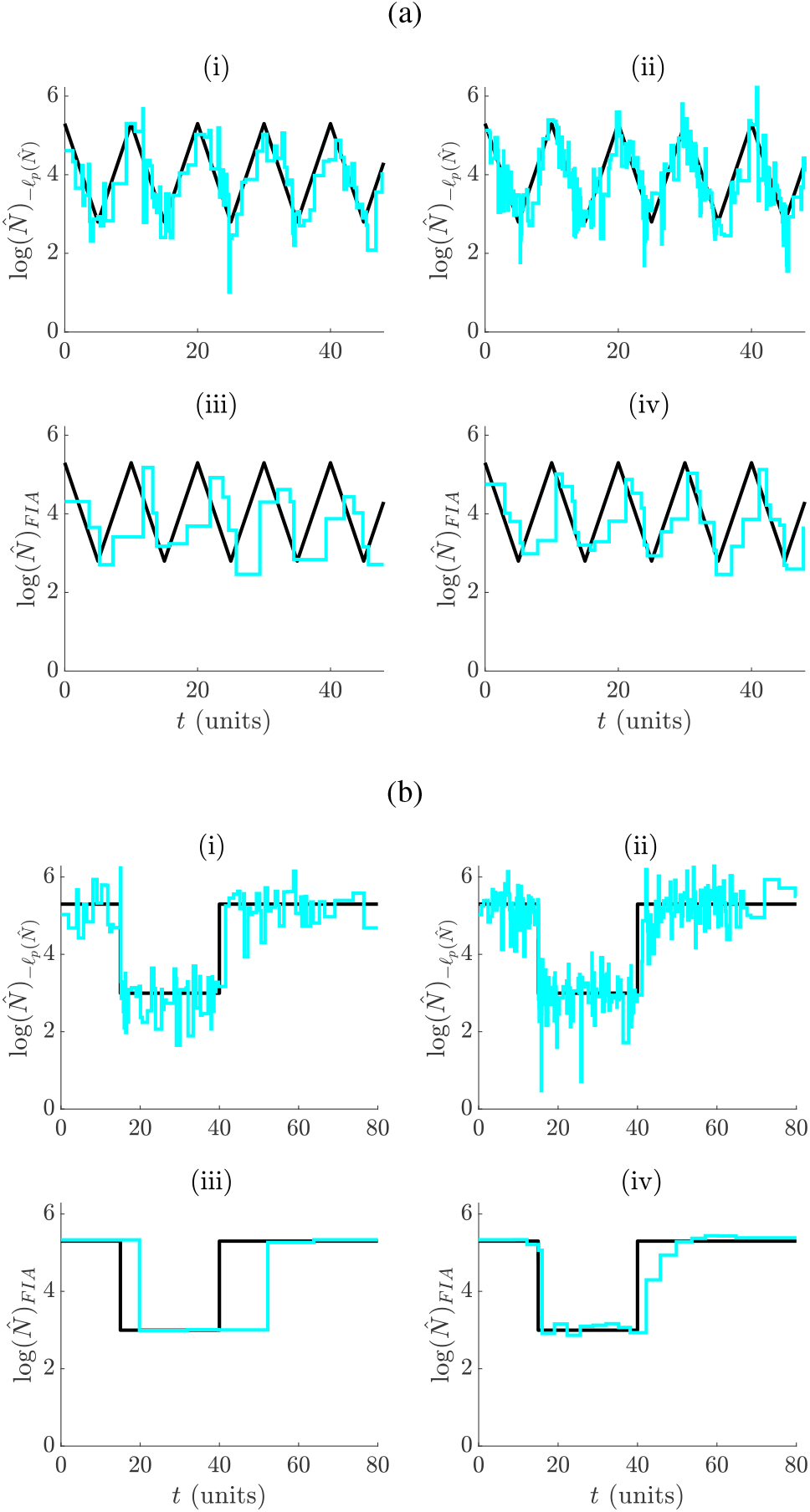
Adaptive periodic and bottleneck estimation with FIA. For (a) and (b), graphs (i)-(ii) present inferred *N*(*t*) under optimal log-likelihood groupings, while (iii)-(iv) show corresponding estimates under the FIA at *ν* = 10^3^. Graphs (i) and (iii) feature *m* = 400 while (ii) and (iv) have *m* = 1000 (data size increases). Panels (a) and (b) respectively consider periodically exponential and bottleneck population size changes, with phylogenies sampled approximately uniformly over [0, 50] and [0, 60] time units.

We consider two model selection problems involving a piecewise-constant *N*(*t*), to formally evaluate the FIA against the QK, BIC and AIC. We slightly abuse notation by redefining *m* as the number of coalescent events per piecewise segment. The first is a binary hypothesis test between a Kingman coalescent null model [16] and an alternative with a single shift to *N*(*t*). We investigate this problem in the Appendix and show in Fig. A2 (i) that the FIA is, on average, better at selecting the true model than other criteria, with the QK a close second. Further, these metrics generally improve in accuracy with increased data. Closer examination also reveals that the FIA and QK have the best overall true positive and lowest false positive rates (Fig. A2(ii)).

The second classification problem is more complex, requiring selection from among 5 possible square waves, with half-periods that are powers of 2. We define 15 change-point times at multiples of *τ* = 50 time units (i.e. there are 16 components) and allow *N*(*t*) to fluctuate between maximum *N*_max_ and ½ *N*_max_. At each change-point and 0, equal numbers of samples are introduced, to allow approximately *m* coalescent events per component (the phylogeny has 16*m* total events). The possible models are in Fig. 6a. A similar problem, but for Gaussian MDL selection, was investigated in [11]. We simulate 200 phylogenies from each wave and compute the probability that each metric selects the correct model (i.e. 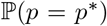) at *N*_max_ = 300 ((i)) and 600 ((ii)) with *ν* = 10^3^ in Fig. 6b. The group size (*k*) search space is *m* times the half-period of every wave.

**Fig. 6:**
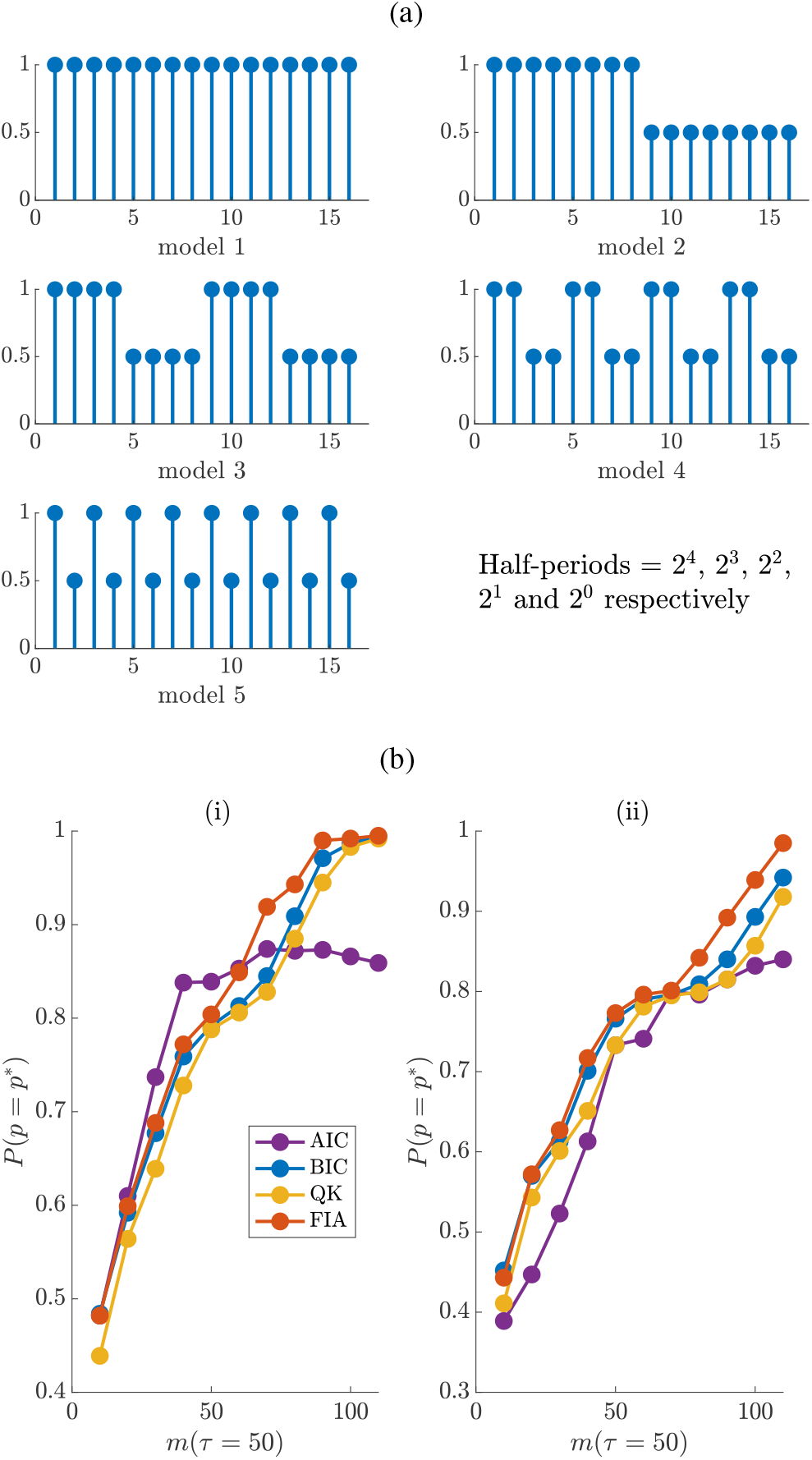
Skyline model selection. We simulate 200 sampled phylogenies from each of the 5 square wave models of (a), with *m* coalescent events per segment. Each square wave varies between *N*_max_ and ½ *N*_max_ (ratios shown on y axes), and occurs with varying half-periods over 16 segments (x axes) of duration *τ*. Each phylogeny contains sampled tips at 0 and every multiple of *τ* time units after. Panel (b) gives the probability that several model selection criteria select the true (*p*\*) model from among these waves at *ν* = 10^3^ for *N*_max_ = 300 ((i)) and *N*_max_ = 600 ((ii)). The FIA is the most accurate criteria on average and improves with *m* and as *ν* gets closer to the true *N*_max_.

The FIA has the best overall accuracy at both *N*_max_ settings, though the BIC is not far behind. The QK displays slightly worse performance than the BIC and the AIC is the worst (except at low *m*). At *N*_max_ = 300 ((i)), there is a greater mismatch with *ν* and so the FIA is not as dominant. As *N*_max_ = 600 ((ii)) gets closer to *ν* this issue dissipates. We discuss this dependence of FIA on *ν* in the Appendix (see Fig. A3). Observe that the 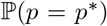 improves for most metrics as the sample phylogeny data size (*m*) increases (consistency). The strong performance of the FIA confirms the impact of parametric complexity, while the suboptimal QK curves suggest that these advantages are sometimes only realisable when this complexity component is properly specified.

## IV. Discussion

Identifying the salient fluctuations in effective population size, *N*(*t*), and effective reproduction number, *R*(*s*), is essential to understanding the retrospective and continuing behaviour of an epidemic, at the population level. A significant swing in *R*(*s*) could inform on whether an outbreak is exponentially growing (e.g. if *R*(*s*) > 1 for a sustained period) or if enacted control measures are working (e.g. if *R*(*s*) falls rapidly below 1) [4] [8]. Similarly, sharp changes in *N*(*t*) could evidence the historical impact of a public health policy (e.g. if *N*(*t*) has a bottleneck or logistic growth) or corroborate hypotheses about past transmissions (e.g. if *N*(*t*) correlates with seasonal changes) [34][31]. Together, *N*(*t*) and *R*(*s*) can provide a holistic view of the temporal dynamics of an epidemic, with their change-points signifying the impact of climatic, ecological and anthropogenic factors [13].

Piecewise-constant approaches, such as the skyline and renewal models, are a tractable and popular way of separating insignificant fluctuations (the constant segments) from meaningful ones (the change-points). However, the efficacy of these models requires principled and data-justified selection of their dimension, *p*. Failure to do so, as in Fig. 2, could result in salient changes being misidentified (i.e. underfitting) or random noise being over-interpreted (i.e. overfitting). Existing approaches to *p*-selection for renewal models usually involve heuristics or trial and error [4]. Skyline models feature a more developed set of *p*-selection methods but many of these, though widely used, are either computationally complex (e.g. involving sophisticated MCMC algorithms) [12] or difficult to interpret (e.g. when *p* is implicitly controlled with smoothing prior distributions) [13] [29].

We therefore focussed on finding a *p*-selection metric that favourably compromises among simplicity, transparency and performance. We started by proving that ascribing *p* solely on the evidence of the log-likelihood (i.e. the model fit) guarantees overfitting (see Eq. (11)). Consequently, it is absolutely necessary to penalise the log-likelihood with a measure of model complexity. However, getting this measure wrong can just as easily lead to underfitting. This is a known issue in common skyline methods that apply smoothing prior distributions for example, where the prior-induced penalty is unclear [20]. Standard metrics, such as the AIC and BIC, are easy to compute and offer transparent penalties; treating model complexity as either equivalent to *p* or *p* mediated by the observed data size (see Eq. (12) and Eq. (13)). However, this description, while useful, is incomplete, and neglects parametric complexity [36].

Parametric complexity describes how the functional relationship among parameters matters. MDL and BMS, which are the most powerful model selection methods, both account for parametric complexity but are often intractable [10]. The general FIA of Eq. (8) approximates both the MDL and BMS and defines this complexity as an integral across parameter space [21]. Unfortunately, this integral is often difficult to evaluate, also rendering the FIA impractical. However, we found that the piecewise-constant nature of renewal and skyline models, together with their Poisson data structures, allowed us to analytically solve this integral and obtain Eq. (14) and Eq. (15). These form our main results, are of similar computability to the AIC and BIC, and disaggregate model complexity into interpretable elements as follows for Eq. (15).

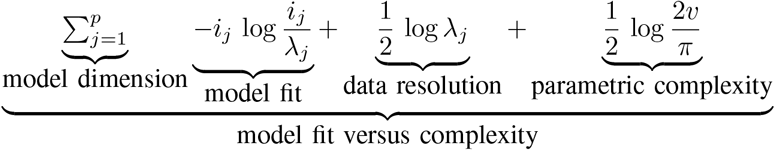

A similar breakdown exists for Eq. (14). Intriguingly, the parametric complexity now only depends on the unknown parameter domain maximum, *ν*.

Knowledge of *ν* is the main cost of our metric. This parameter limit requirement is not unusual and can often improve estimates. In [26] and [27] this knowledge facilitated exact inference from sampled phylogenies, for example. Similar domain choices are also implicitly made when setting prior distributions on *R*(*s*) and *N*(*t*) or practically performing MCMC sampling. In Fig. A3 we explored the effect of misspecifying *ν*. While drastic mismatches between the true and assumed *ν* can be detrimental, we found that in some cases poor knowledge of *ν* can be inconsequential. We adapted the QK metric [33] to obtain Eq. (16) and Eq. (17) which, though less interpretable than the FIA, also somewhat account for parametric complexity and offer good performance should reasonable knowledge of *ν* be unavailable.

The FIA balances performance with simplicity. The MDL method it approximates has the desirable theoretical properties of generalisability (it mediates overfitting and underfitting) and consistency (it selects the true model with increasing probability as data accumulate) [10]. We therefore investigated whether the FIA maintained these properties. In Fig. 3 and Fig. 5 we demonstrated that the FIA not only inherits the generalisability property, but also regulates its selections based on the available data. Higher data resolution supports larger *p* as both bias and variance can be simultaneously reduced under these conditions [41]. Fig. 4, Fig. 6, Fig. A1 and Fig. A2 confirmed the consistency of the FIA, in addition to benchmarking its performance against the comparable AIC and BIC. We found that the FIA consistently outperformed all other metrics, provided that *ν* was not drastically misspecified.

We recommend the FIA as a principled, transparent and computationally simple means of adaptively estimating informative changes in *N*(*t*) and *R*(*s*), and for diagnosing the relative contributions of different components of model complexity. Specifically, the FIA can be easily interfaced with the EpiEstim and projections packages [4] [23], which are common renewal model toolboxes for analysing real epidemic data, to formalise the window size choices used in *R*(*s*) inference. Until now, these choices have been subjective. For skyline analyses, we propose the FIA as a useful diagnostic for verifying the *N*(*t*) estimates generated by phylogenetic software such as BEAST or phylodyn [39] [14]. This can help validate or interrogate the outputs of common but complex MCMC methods and guard against known issues such as oversmoothing (underfitting) [20].

Sampled phylogenies and incidence curves, and hence skyline and renewal models, have often been treated separately in the epidemiological and phylodynamics literature. While they do solve different problems, we showed how refocussing on their shared piecewise-Poisson framework exposed their common complexity properties. Our information theoretic approach could also generate broad insight into other distinct models in genetics, molecular evolution and ecology [28]. The structured coalescent model is often used to estimate migration rate and population size changes from phylogeographic data [2] while sequential Markovian coalescent methods are widely applied to infer demographic changes from metazoan genomes [19]. These models involve Poisson count and histogram records and piecewise parameter sets and are promising candidates for future application of our metrics.

## Appendix

## Binary Model Selection

We examine the binary classification performance of the FIA, QK, BIC and AIC for both renewal and skyline models. For the first we set *m* = 200 days and use a constant null model with *R*(*s*) = 1.5, to exemplify an uncontrolled epidemic. The alternative model changes to *R*(*s* ≥ *m*/2) = 0.5, simulating rapid control at *m*/2 (inset of Fig. A1). We randomly generate 10^3^ epidemics with some null model probability (*P* (*p*\* = 1)) and compute the frequentist probability that each criteria selects the correct model (*P* (*p* = *p*\*)) in Fig. A1. We find that the FIA uniformly outperforms all other criteria, with the QK as its closest competitor. The AIC performs poorly, as does 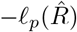 (not shown), because they are biased towards the more complex model. Relative metric performance is unchanged if we instead set *R*(*s ≥ m*/2) = 2.5 (an accelerating epidemic).

**Fig. A1:**
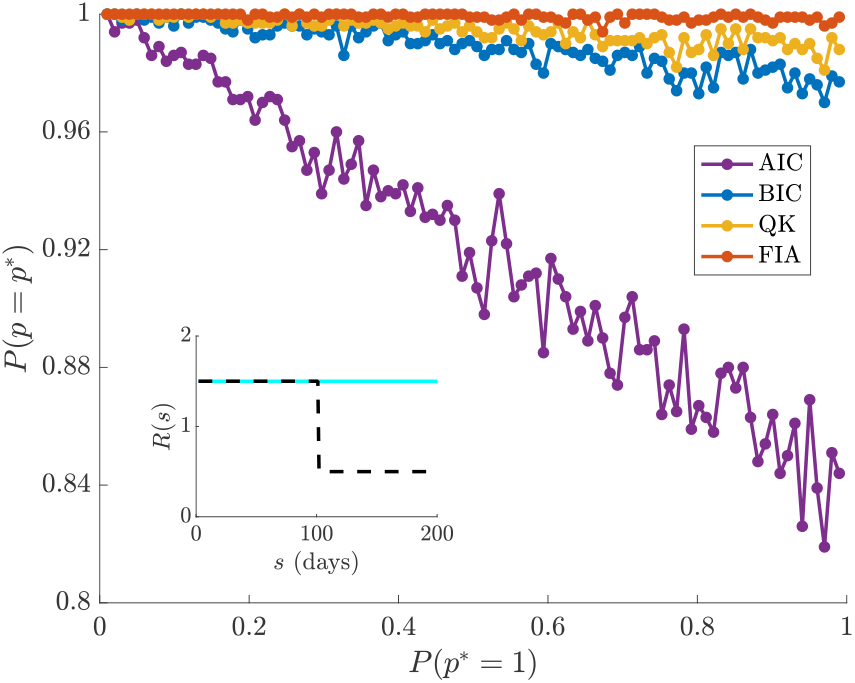
Binary renewal model selection. The consistency of several selection criteria is tested on a binary classification problem in which the null model 1 has no change in *R*(*s*) (solid, inset), while the alternative model 2 has a rapid decline (dashed, inset). We generate 10^3^ independent incidence curves randomly according to model 1 with probability 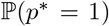, and compute the ability of each criteria to decipher the correct model, 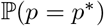. The FIA outperforms other metrics at every *p*\* with QK a close second.

For the skyline problem we test between a Kingman coalescent null model [16] with *N*_1_ = 1000, and an alternative with a single shift to *N*_2_ = 500 that simulates rapid change potentially due to some environmental driver at *τ* = 250 units. We set *ν* = 10^5^ and generate 500 replicate phylogenies, with *m* controlling the quantity of data available per piecewise component (so the total number of coalescent events is 2*m*). This is a slight abuse of previous definitions of *m* but is more useful here as we want *p*\* = 1 for the null model and *p*\* = 2 for the alternative. We introduce sampled tips at 0 and *τ* time units only. The grouping parameter search space is 4 ≤ *k* ≤ 2 *m* with *p* = 2*m*/*k*. Fig. A2 presents our main results, showing that the FIA is, overall, more accurate (achieving a higher 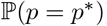) with the QK second. We find relative performance is largely unchanged with *ν* and when *τ* is doubled. Observe that all metrics except the AIC (which is known to be inconsistent) improve with data size, *m*.

## Weaknesses of Piecewise Model Selection

In Results we found the FIA to be a viable and top performing model selection strategy, when compared to standard metrics of similar computability such as the AIC and BIC. However, the FIA can do worse if the parameter maximum *ν* is large relative to the actual space from which *R*(*s*) or *N*(*t*) is drawn. In such cases, the incorrect parameter bounds can cause the FIA to overestimate the complexity of the generating renewal or skyline models. While the QK criteria offers a more stable and reasonably performing MDL alternative, it is less interpretable. Here we examine the nature of this *ν* dependence, and discuss some general issues limiting piecewise model selection.

**Fig. A2:**
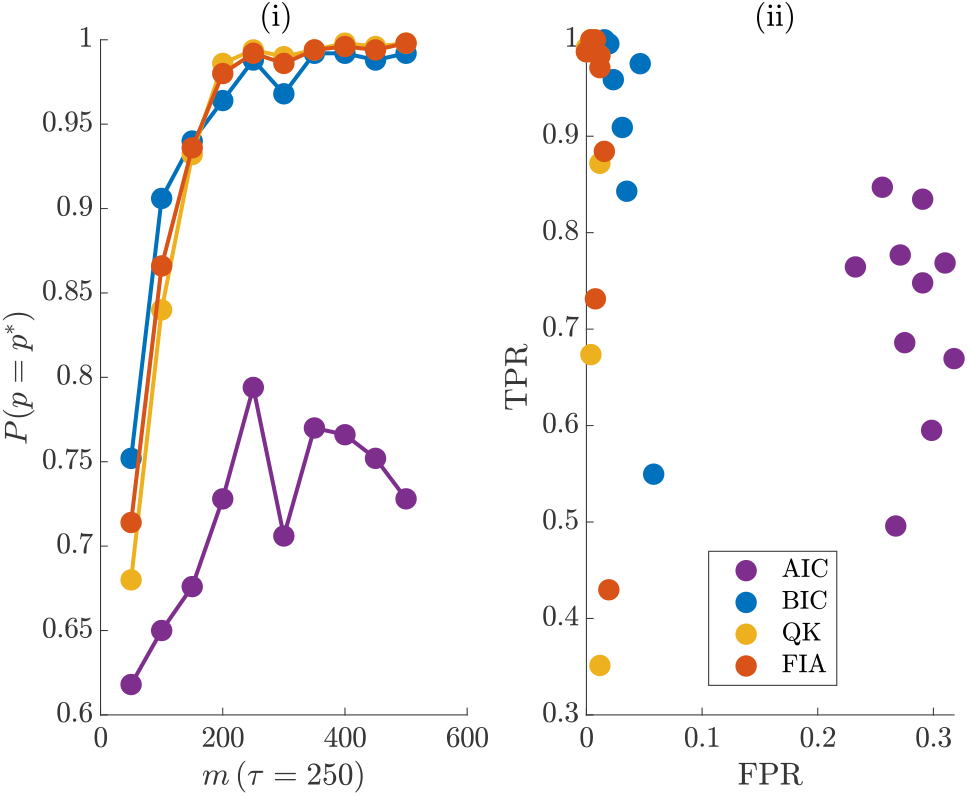
Binary skyline model selection. We simulate 500 conditionally independent phylogenies from skyline models and test the classification ability of model selection criteria. The null model is a Kingman coalescent with *N*_1_ = 1000, and the alternative features a sharp fall to *N*_2_ = 500 at *τ* = 250 time units. The sampled tips of the phylogeny are introduced at 0 and *τ* only. Graph (i) gives the probability of correct classification 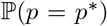 as a function of data size *m*. The FIA performs best, on average, but the BIC is better at small *m*. Graph (ii) gives the true (TPR) and false positive rates (FPR). The FIA and QK have the best overall rates.

In Fig. 4b(ii) we showed the FIA outperforming other metrics for a model selection problem over piecewise *R*(*s*) functions drawn within the artificial range [0.5, 1.5] (the AIC was better at higher *p* due to its tendency to overfit). We achieved this by setting *ν* to the true *R*_max_ = 1.5. However, when there is a significant mismatch between *ν* and *R*_max_ we find that the FIA is notably inferior to the QK and BIC. Fig. A3a illustrates, at *ν* = 100 and 6, how the magnitude of this mismatch influences relative performance. However, this effect is not always important, as seen in Fig. 4b(i) where *R*_max_ = 6 and *ν* = 100. The skyline model also has this FIA *ν*-dependence. We re-examine the square wave model selection problem of Fig. 6b, but for *ν* ranging between 10^2^ and 10^5^. Fig. A3b plots the resulting changes in the FIA detection probability at *N*_max_ = 300 ((i)) and 600 ((ii)). There we observe, that while the FIA is sensitive to *ν*, it still performs well over the entire range. Thus, the FIA can sometimes be a choice selection metric, even in the absence of reasonable parameter space knowledge.

**Fig. A3:**
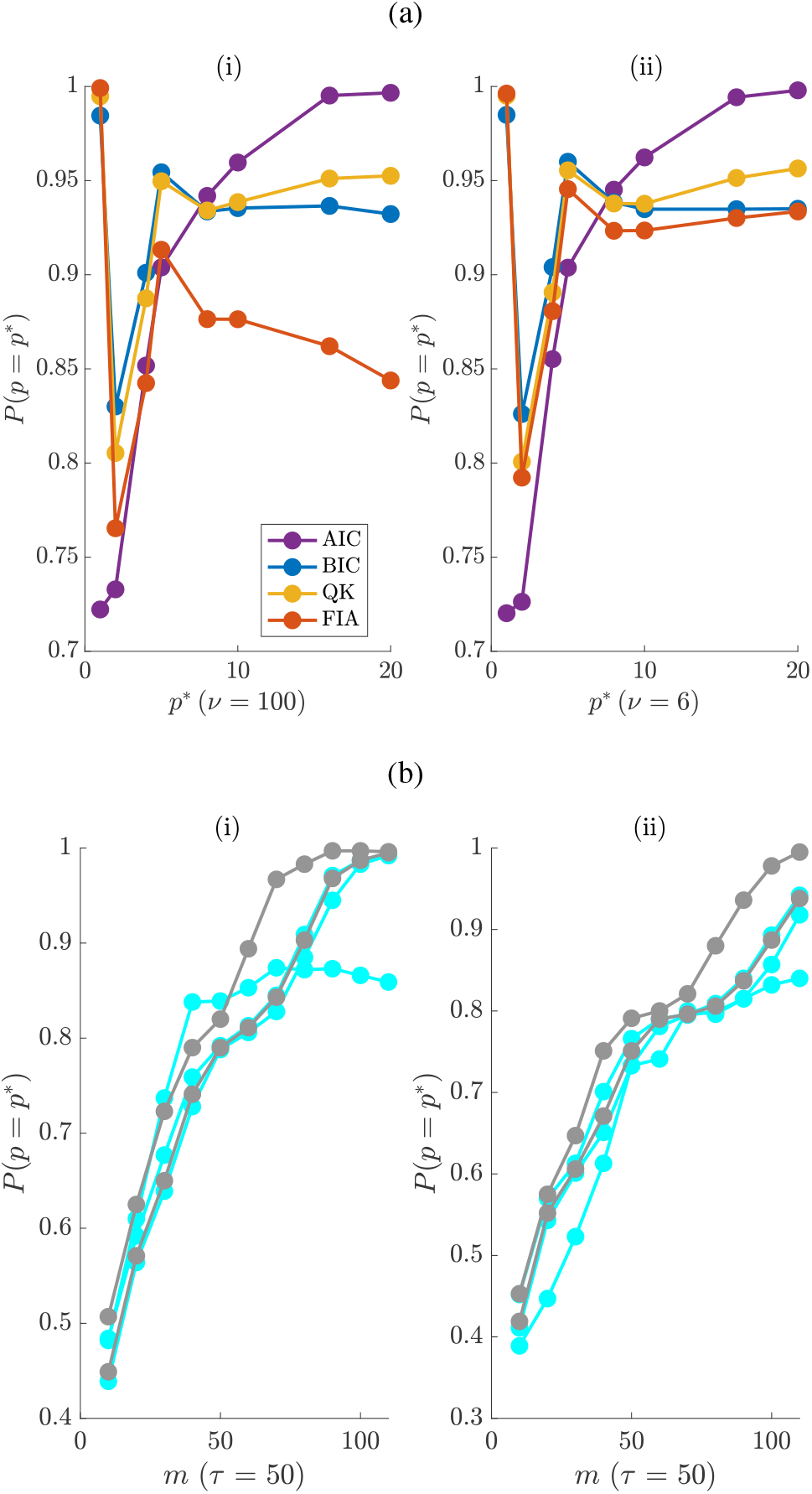
FIA parameter space sensitivity. In (a) we repeat the simulations from Fig. 4b(ii) but at different *ν*. The accuracy of the FIA clearly depends on the discrepancy between *ν* and *R*_max_ = 1.5, and becomes inferior when *ν* is dramatically above this maximum ((i)). In (b) we revisit the simulations of Fig. 6a, but vary *ν* between 10^2^ and 10^5^. The AIC, BIC and QK from Fig. 6a are in cyan, while the best and worst case FIA values are in grey. While the FIA does depend on *v*, interestingly, its performance is still superior on average. for both *N*_max_ = 300 ((i)) and 600 ((ii)).

Lastly, we comment on some general issues limiting *p*-selection performance of any metric on renewal and skyline models. The MLEs and FIs of the renewal model depend on the *λ*_*j*_ and *i*_*j*_ groups. As a result, epidemics with low observed incidence (i.e. likely to have *i*_*j*_ = 0) and diseases possessing sharp (low variance) generation time distributions (i.e. likely to feature *λ*_*j*_ = 0) will be difficult to adaptively estimate. Hence, we conditioned on the outbreak not dying out. Similarly, the MLEs and FIs of the skyline are sensitive to *m*_*j*_, meaning that it is necessary to ensure each group has coalescent events falling within its duration. Forcing segment end-points to coincide with coalescent events, as in [6], guards against this identifiability problem [28]. However, skyline model selection remains difficult even after averting this issue.

This follows from the random timing of coalescent events, which means that regular *k* groupings can miss change-points, and that long branches can bias analysis [25]. These are known skyline plot issues and evidence why we truncated the last few events in the *N*(*t*) simulations. Further, there will always be limits to the maximum temporal precision attainable by *R*(*s*) and *N*(*t*) estimates under renewal and skyline models. It is impossible to infer changes in *R*(*s*) on a finer time scale than that of the observed incidence curve or estimate more *N*(*t*) segments than the number of available coalescent events [28]. This cautions against naively applying the criteria we have developed here. It is necessary to first understand and then prepare for these preconditions before sensible model selection results can be obtained. Practical model selection is rarely straightforward and the performance of most metrics is often only strictly guaranteed asymptotically [10].

## Empirical Case Study: HIV-1

We consider an empirical, ultrametric phylogeny composed of HIV-1 sequences sampled in 1997 from the Democratic Republic of Congo. This dataset was previously examined in [38]. In Fig. A4 we illustrate several estimates of the effective population size underling this phylogeny. As a baseline we plot the classic skyline plot [32] for this phylogeny in grey on (i), (ii) and (iv). This represents the maximally parametrised skyline model and is known to overfit. Because the classic skyline converts every coalescent interval into a population size estimate, it also portrays where the events in the HIV tree are located. The clustering of estimates across 1940-1980 indicates that this period is notably more informative (i.e. has a higher count record event density) than others around it.

We investigate two extremes of skyline model selection methodology. In Fig. A4(i) we consider the generalised skyline plot, which uses a small sample AIC. This method is simple, computable and improves on the noisy classic skyline by using *p*\* = 13. However, it does require more extensive optimisation than our metrics (it chooses groups based on their durations) and can be susceptible to overfitting [15]. In (ii) we plot estimates from the multiple change-point method of [24]. This approach is computationally intensive and lacks transparency but uses powerful reversible jump MCMC algorithms. Its output smooths over all demographic fluctuations.

In (iv) we compute the FIA solution with *ν* = 2000, which mediates between (i) and (ii). The FIA responds to the varying data-density across the tree by using notably more parameters than (i) in the 1940-1980 period, where it can be confident of a smooth trend, and fewer otherwise. This approach to group choice was theoretically supported in [28]. In (iii) we compare the FIA (*p*\* = 16) and BIC (*p*\* = 8) curves, where we find that they agree at small p due to the large sample size in that region (the BIC is an asymptotic approximation to the FIA). However, at larger p, where the space of parameter interactions is more notable, the parametric complexity terms matter.

**Fig. A4:**
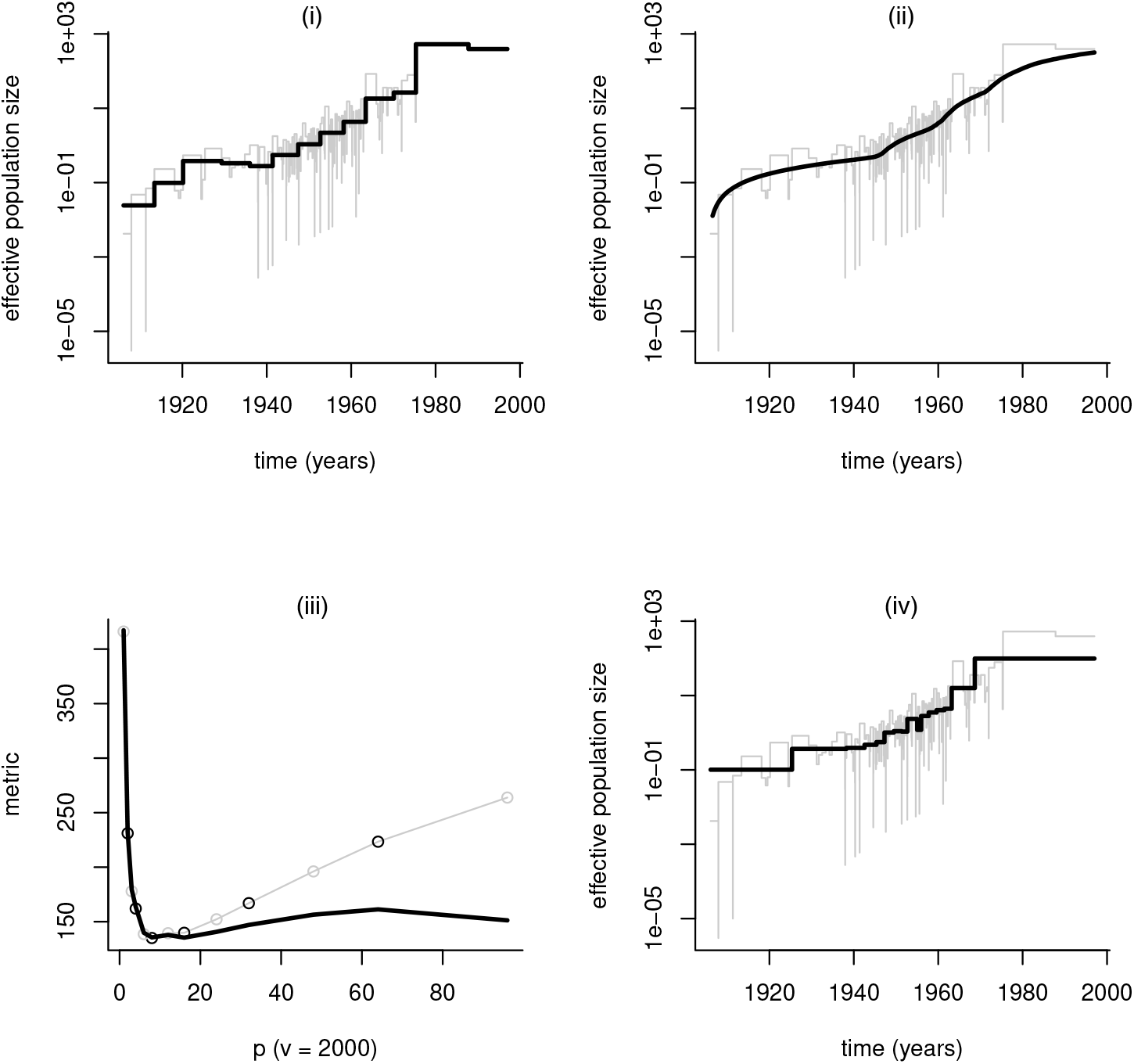
HIV demographic estimates. We estimate the population size history underlying an empirical HIV phylogeny with *m* = 192 coalescent events. All tips of this tree are sampled in 1997 from the Democratic Republic of Congo. We plot the generalised skyline, the multiple change-point method and the FIA-optimised skyline estimates in (i), (ii) and (iv) in black against the classic skyline in grey. In (iii) we show the optimisation over p of the FIA (black) against its associated BIC (grey).

## References

[1] A Barron, J Rissanen, and B Yu. The minimum description length principle in coding and modeling. IEEE Trans. Info. Theo, 44(6):2743–60, 1998.

[2] P Beerli and J Felsenstein. Maximum Likelihood Estimation of a Migration Matrix and Effective Population Sizes in n Subpopulations by using a Coalescent Approach. PNAS, 98(8):4563–68, 2001.

[3] T Churcher, J Cohen, J Novotny, et al. Measuring the path toward malaria elimination. Science, 344(6189):1230–2, 2014.

[4] A Cori, N Ferguson, C Fraser, et al. A new framework and software to estimate time-varying reproduction numbers during epidemics. Am. J. Epidemiol, 178(9):1505–12, 2013.

[5] T Cover and J Thomas. Elements of Information Theory. John Wiley and Sons, second edition, 2006.

[6] A Drummond, A Rambaut, B Shapiro, and O Pybus. Bayesian coalescent inference of past population dynamics from molecular sequences. Mol. Biol. Evol, 22(5):1185–92, 2005.

[7] C Fraser. Estimating individual and household reproduction numbers in an emerging epidemic. PLoS One, 8:e758, 2007.

[8] C Fraser, D Cummings, D Klinkenberg, et al. Influenza transmission in households during the 1918 pandemic. Am. J. Epidemiol, 174(5):505–14, 2011.

[9] M Gill, P Lemey, N Faria, A Rambaut, B Shapiro, and M Suchard. Improving Bayesian Population Dynamics Inference: A Coalescent-Based Model for Multiple Loci. Mol. Biol. Evol, 30(3):713–24, 2012.

[10] P Grunwald. The Minimum Description Length Principle. The MIT Press, 2007.

[11] A Hanson and P Fu. Advances in Minimum Description Length: Theory and Applications, chapter Applications of MDL to selected families of models. MIT Press, 2004.

[12] J Heled and A Drummond. Bayesian inference of population size history from multiple loci. BMC Evol. Biol, 8(289), 2008.

[13] S Ho and B Shapiro. Skyline-plot methods for estimating demographic history from nucleotide sequences. Mol. Ecol. Res, 11:423–34, 2011.

[14] M Karcher, J Palacios, S Lan, et al. PHYLODYN: an R package for phylodynamic simulation and inference. Mol. Ecol. Res, 17:96–100, 2017.

[15] R Kass and A Raftery. Bayes factors. J. Amer. Stat. Assoc, 90(430):773–95, 1995.

[16] J Kingman. On the genealogy of large populations. J. Appl. Prob, 19:27–43, 1982.

[17] E Lehmann and G Casella. Theory of Point Estimation. Springer-Verlag, second edition, 1998.

[18] P Lemey, O Pybus, B Wang, et al. Tracing the origin and history of the HIV-2 epidemic. PNAS, 100(11):6588–92, 2003.

[19] H Li and R Durbin. Inference of human population history from individual whole-genome sequences. Nature, 475(7357):493–6, 2011.

[20] V Minin, E Bloomquist, and M Suchard. Smooth skyride through a rough skyline: Bayesian coalescent-based inference of population dynamics. Mol. Biol. Evol, 25(7):1459–71, 2008.

[21] J Myung, D Navarro, and M Pitt. Model selection by normalized maximum likelihood. J. Math. Psychol, 50:167–79, 2006.

[22] M Nordborg. Handbook of Statistical Genetics: Coalescent Theory. John Wiley and Sons, 2001.

[23] P Nouvellet, A Cori, T Garske, et al. A simple approach to measure transmissibility and forecast incidence. Epidemics, 22:29–35, 2018.

[24] R Opgen-Rhein, L Fahrmeir, and K Strimmer. Inference of demographic history from genealogical trees using reversible jump Markov chain Monte Carlo. BMC Evol. Biol, 5(6), 2005.

[25] K Parag, L du Plessis, and O Pybus. Jointly inferring the dynamics of population size and sampling intensity from molecular sequences. Mol. Biol. Evol, msaa016, 2020.

[26] K Parag and O Pybus. Optimal point process filtering and estimation of the coalescent process. J. Theor. Biol, 421:153–67, 2017.

[27] K Parag and O Pybus. Exact bayesian inference for phylogenetic birth-death models. Bioinformatics, 34(21):3638–45, 2018.

[28] K Parag and O Pybus. Robust design for coalescent model inference. Syst. Biol, 68(5):730–43, 2019.

[29] K Parag, O Pybus, and C Wu. Are skyline plot-based demographic estimates overly dependent on smoothing prior assumptions? BioRxiv, 920215, 2020.

[30] M Pitt, I Myung, and S Zhang. Toward a method of selecting among computational models of cognition. Psych. Rev, 109(3):472–91, 2002.

[31] O Pybus, M Charleston, S Gupta, et al. The epidemic behavior of the hepatitis C virus. Science, 292(5525):2323–5, 2001.

[32] O Pybus, A Rambaut, and P Harvey. An integrated framework for the inference of viral population history from reconstructed genealogies. Genetics, 155:1429–37, 2000.

[33] G Qian and H Kunsch. Some notes on Rissanen’s stochastic complexity. IEEE Trans. Info. Theo, 44(2):782–6, 1998.

[34] A Rambaut, O Pybus, M Nelson, et al. The genomic and epidemiological dynamics of human influenza A virus. Nature, 453(7195):615–619, 2008.

[35] J Rissanen. Modeling by shortest data description. Automatica, 14:465–71, 1978.

[36] J Rissanen. Fisher information and stochastic complexity. IEEE Trans. Info. Theo, 42(1):40–7, 1996.

[37] D Snyder and M Miller. Random Point Processes in Time and Space. Springer-Verlag, 2 edition, 1991.

[38] K Strimmer and O Pybus. Exploring the demographic history of DNA sequences using the generalized skyline plot. Mol. Biol. Evol, 18(12):2298–305, 2001.

[39] M Suchard, P Lemey, G Baele, et al. Bayesian phylogenetic and phylodynamic data integration using BEAST 1.10. Virus Evol, 4(vey016), 2018.

[40] P Turchin. Complex Population Dynamics: A Theoretical/Empirical Synthesis. Princeton University Press, 2003.

[41] T van Erven and P Grunwald. Catching up faster by switching sooner: a predictive approach to adaptive estimation with an application to the AIC–BIC dilemma. J. R. Statist. Soc. B, 74(3):361–417, 2012.

[42] J Wallinga and M Lipsitch. How generation intervals shape the relationship between growth rates and reproductive numbers. Proc. R. Soc. B, 274:599–604, 2007.

[43] J Wallinga and P Teunis. Different epidemic curves for severe acute respiratory syndrome reveal similar impacts of control measures. Am. J. Epidemiol, 160(6):509–16, 2004.

